# Exploring methanogenic archaea and their thermal responses in the glacier-fed stream sediments of Rongbuk River Basin, Mt. Everest

**DOI:** 10.1101/2024.11.15.623738

**Authors:** Wei Ma, Miao Lin, Peihua Shen, Hongfei Chi, Weizhen Zhang, Jingyi Zhu, Shaoyi Tian, Pengfei Liu

## Abstract

Glacier-fed streams (GFS) are emergent sources of greenhouse gas methane, and methanogenic archaea in sediments contribute largely to stream methane emissions. However, little is known about the methanogenic communities in GFS sediments and their key environmental driving factors. This study analyzed stream sediments from the Rongbuk River basin on Mount Everest for methanogenic communities and their temperature responses through anaerobic microcosm incubations at 5°C and 15°C. Diverse methanogens were identified, including acetoclastic, hydrogenotrophic, and methylotrophic types. Substantial methane and CO_2_ production were detected across altitudes and increased significantly at 15°C, with both methane and CO_2_ production rate negatively correlated with altitude. While temperature sensitivity of CO_2_ production but not CH_4_ showed a negative altitude correlation. Methanogens increased substantially over long-term incubation, dominating the archaeal community. At 15°C, the relative abundance of several methanogenic groups was strongly correlated with altitude, with positive correlations observed for *Methanomassiliicoccaceae* and *Methanoregulaceae*, and negative correlations for *Methanocellaceae* and *Methanotrichaceae*, respectively. Besides altitude, factors like phosphorus, C-to-N ratio, and pH also affected methanogenic structure, methane and CO_2_ production, and temperature sensitivities. This study offers new insights into methanogens and methane production in GFS sediments, improving our understanding of GFS carbon cycling and its potential responses to climate change.

## Introduction

Methane is the second most important anthropogenic greenhouse gas after CO_2_, accounting for about 25% of the observed warming to date and playing a major role in controlling the Earth’s climate (Conrad 2009, Saunois et al. 2020). Atmospheric CH_4_ concentrations have nearly tripled since the Industrial Revolution, increasing from 720 ppbv before 1750 to 1930 ppbv in 2024 (NOAA). Rivers and streams are recognized as an important source of CH_4_(Battin et al. 2023, Li et al. 2021, Song et al. 2024, Stanley et al. 2016, Zhang et al. 2020b), with an annual global emission of ∼27.9 Tg CH_4_ per year, which is roughly equal in magnitude to those of other freshwater systems (Rocher-Ros et al. 2023). Globally, headwater streams are significant sources of CH_4_, which contribute to 75.5% of the global riverine CH_4_ emissions(Li et al. 2021). Glacier-fed streams (GFS) are special types of headwater streams and constitute a prominent geomorphological and ecological component of the glacier foreland (Wilhelm et al. 2013), which are significantly affected by climate change and glacier melting (Busi et al. 2022, Milner et al. 2017). Recent field observation studies have revealed that GFS is a new hotspot of CH_4_ emission (Du et al. 2024, Gao et al. 2023, Konya et al. 2024, Lamarche-Gagnon et al. 2019b, Zhang et al. 2021a). Moreover, with rising temperatures, accelerated glacier melting, and increased glacier runoff under climate change, methane emissions from GFS may increase in the future(Gao et al. 2023, Zhang et al. 2021a). Therefore, revealing the methane emission process in GFS is of great significance to understand the carbon cycling of alpine ecosystems and its response and feedback to climate change.

Methanogenic archaea (methanogens) are obligate anaerobic microorganisms (Liu and Whitman 2008). They are the key microbial groups that drive methane production in water-logged anaerobic ecosystems such as lake sediments and wetlands (Conrad 2009, Conrad 2020). The role of methanogens and their contribution to methane production in river sediments have been traditionally neglected since the river sediments are well-aerated(Stanley et al. 2016). However, studies in recent years have shown that the methane emitted from streams is mainly produced in sediments by methanogens (Bodmer et al. 2020, Comer-Warner et al. 2018). Observed methane production potential is comparable to that of freshwater lake sediments (Bednarik et al. 2017, Bednařík et al. 2019, Bodmer et al. 2020, Crawford et al. 2017, Wilkinson et al. 2019, Wilkinson et al. 2015). Abundant and diverse methanogens live in rivers and streams sediments, with the number of methanogens in a range of 10^5^-10^8^ (Bednařík et al. 2019, Brablcová et al. 2015, Chaudhary et al. 2017). Three main types of methanogens, including hydrogenotrophic, acetoclastic and methylotrophic methanogens, are well detected (Bednařík et al. 2019, Brablcová et al. 2015, Chaudhary et al. 2017, Nagler et al. 2021). These methanogens may live in anoxic and hypoxic pockets (“anaerobic microzones”) of the well-oxygenated hyporheic zone(Battin et al. 2016, Kohler et al. 2024, Romeijn et al. 2019). In addition, many methanogens have an amazing ability to tolerate a certain amount of oxygen(Angle et al. 2017, Conrad et al. 2014), which might enable their survival in river sediment. Compared with investigated rivers and streams, the water flow of GFS is more intense, leading to a more severe disturbance and aeration(Michoud et al. 2023, Wilhelm et al. 2013). GFS has obvious seasonal changes in water levels due to glacier discharge, so their sediments are in frequent aerobic-anaerobic alternations. Moreover, GFS is an extreme environment with low temperatures, strong radiation, and poor substrates(Busi et al. 2022, Milner et al. 2017). Currently, there was only one study that identified the presence of methanogens in the GFS sediment biofilms of Mount Stanley, Africa(Michoud et al. 2023). Therefore, the diversity of the methanogens and methane production processes in the sediments of GFS are largely unknown.

In addition, GFS usually have large altitude differences within a short range, and the properties of the streams are affected by the distance from the glacier. For example, water temperature, nutrients, etc. Existing studies have shown that microbial community structure and diversity have obvious altitude gradient distribution characteristics (Battin et al. 2016, Wilhelm et al. 2015, Wilhelm et al. 2013). However, it is not clear whether the distribution of methanogens community in GFS sediments has an altitude gradient. Temperature is an important factor affecting methanogens (Conrad 2023, Yvon-Durocher et al. 2014). Studies have shown that the respiration and biological methane production processes have a significant response to temperature(Acuna et al. 2008, Comer-Warner et al. 2018). The increase in temperature with climate change is amplified at high-altitude ecosystems under climate change. However, our understanding on the responses of methane production processes in GFS to temperature is still lacking.

Therefore, this study aims to answer the following questions: 1. Are there abundant methanogens with methane production potential in GFS sediments? 2. What are the key factors affecting the methanogenes community in GFS sediments? 3. How will methane production in GFS sediment respond to increased temperatures? We chose the GFS in Rongbuk River Basin of Mount Everest, which relies on melting water of Rongbuk glacier as its main water sources, as our research model (Fig. 1). Along the GFS, sediment samples were collected from 4,369 m to 5,145 m, and anaerobic microcosm incubations of sediments at 5°C and 15°C were carried out to quantify methane production potential and temperature sensitivity. The results revealed that the GFS sediments have obvious methane production activity and diverse methanogens, which were significantly associated with altitude.

**Fig. 1.**
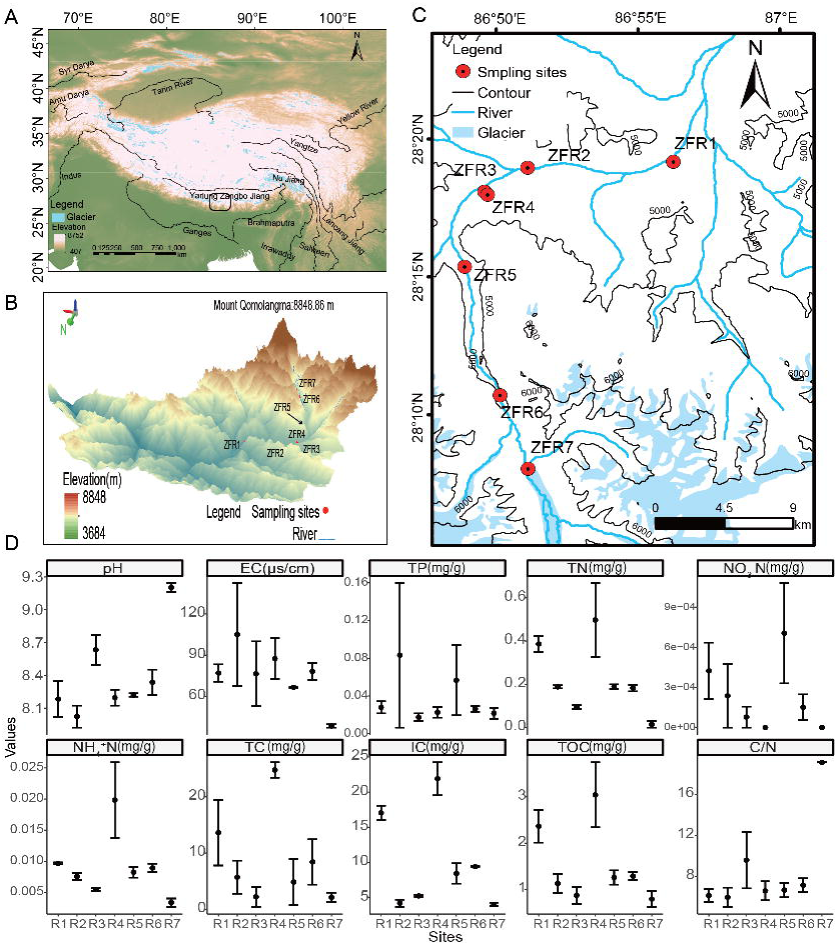
Study area and sampling sites of GFS sediments from the Rongbuk river Basin and major characteristics of the stream sediments. a. the location of the Rongbuk river Basin. b. Distribution of sampling sites along the Rongbuk river Basin, highlighting their distances to the Rongbuk glacier. c. Sampling sites along the Rongbuk Basin, highlighting the altitude distribution of sampling sites. d. major characteristics of the GFS sediments. Data shown are means ± SD, n=3.

## Materials and Methods

### Field Sampling

The Rongbuk River, originating from the Rongbuk Glacier (86.92°E, 27.98°N), is located in the Rongbuk river basin, a typical glacier-fed basin on the northern slope of Mt. Everest, south of the Tibetan Plateau (Fig. 1)(Li et al. 2023). As the largest glacier-fed river on the north slope of Mt. Everest, approximately 70% of the Rongbuk River’s flow is derived from glacier meltwater, with the remaining 30% from precipitation(Li et al. 2023, Ye et al. 2022). Rapid warming in the Mount Everest region has caused a continuous increase in the Rongbuk Glacier’s melting rate over recent decades(King et al. 2020, Ye et al. 2022), with peak meltwater runoff from the Rongbuk River expected around the middle of this century(Sun et al. 2022).

On May 1st–2nd, 2022, we collected stream sediment samples from seven sites along the Rongbuk River (Figs. 1 and S1; Table S1). Sediments were collected using sterile shovels and transferred into 50 mL Falcon tubes and 500 mL Nalgene bottles (Thermo Fisher Scientific, US). During sampling, water temperatures ranged from 4.1°C to 8.4°C, with an average of 5.4°C. After collection, Falcon tubes were stored at −20°C for molecular analysis. The 500 mL Nalgene bottles, filled with sediment, were tightly sealed and stored at 4°C for use in anaerobic microsome incubation and sediment physicochemical property measurements, which were conducted within two months of collection.

### Characterization of sediment physico-chemical properties

Sediment physico-chemical properties were characterized as previously described (Liu et al. 2017a, Liu et al. 2013). Sediment water content was determined by oven drying 10 g of fresh sediments at 105°C for 48 h. The pH and conductivity values were measured after mixing wet soil with distilled water at a soil-to-water ratio of 1:5 (g/g), by the potentiometric method. The total amount of soluble salts was determined by the gravimetric method, as described by(Liu et al. 2017a) (Liu et al. 2016a, Liu et al. 2017b). The total nitrogen content (TN) was determined using the Kjeldahl method(Wang and Oien 1986). Sediments ammonium nitrogen (NH_4_^+^) and nitrate-nitrogen (NO_3_^-^) were extracted from wet soils with 2 M KCl (soil to solution ratio = 1:5) using Smartchem200 Discrete Auto Analyzer (Skalar, Breda, and Netherlands). Total phosphorus (TP) was determined using Smartchem200 Discrete Auto Analyzer. Total potassium (TK) was determined using Atomic Absorption Spectrophotometer (Shimadzu Corp, Japan). The available phosphorus contents (AP) were determined using the acid digestion method. Total carbon (TC) and total organic carbon (TOC) were determined on a milled sample by combustion at 990°C using the TOC Analyzer (Shimadzu Corp, Japan).

### Anaerobic incubation of sediment and chemical analyses

We conducted anaerobic incubations on sediments collected from different altitudes in the Rongbuk River basin, Mount Everest, as follows (Liu et al. 2013):

Briefly, about ∼5 g aliquots [about 3.5 g dry weight (gdw)] of the sediment were transferred in five replicates to 26 mL sterile pressure tubes. The exact amount of sediment was determined gravimetrically. Each tube was then supplemented with 10 mL of sterile water using a pipette, sealed with a butyl rubber stopper and aluminum cap, and purged with high-purity nitrogen gas at 0.05 MPa for 10 minutes to establish an anaerobic environment. Incubations were set up at temperatures of 5°C and 15°C. The anaerobic tubes were incubated at temperatures of 5°C and 15°C in dark without shaking. Gas samples were periodically collected every 10-14 days from the headspace of the tubes using a gas-tight pressure lock syringe (Dynatech), and methane and carbon dioxide concentrations were then measured with an Agilent 8890 gas chromatograph systems using a flame ionization detector after separation at HayeSep Q (2.44 m length, 2 mm diameter) with pure H_2_ and pure N_2_ as carrier gas. After about 281 days and 146 days of incubation at 5°C and 15°C, respectively, triplicate tubes were sacrificed for collecting liquid and sediments. Briefly, each incubation tube was uncapped, and the slurry was decanted into a 15 mL falcon tube and centrifuged for 12 min at 12,000 g. 1.5 mL clarified supernatant were taken and used immediately to measure pH as described above. The remaining liquid was discarded, and the pellet was immediately stored at −80°C for nucleic acid extraction for DNA. At the end of incubation, substantial amounts of CH_4_ and CO_2_ were detected in all samples except for those from site R7 (5,145 m). At R7, only trace amounts of CH_4_ (∼113.3 and 3,990.7 ppm 5°C and 15°C, respectively) were accumulated in the headspace. Due to the limited amount of gas production at R7, the data from this site were not shown, and they were also excluded for further analysis (e.g., Q10 calculations).

### DNA extraction, amplicon sequencing of archaea 16S rRNA gene and data processing

To investigate the specific groups of archaea involved in methane production, we analysed archaeal community structures using 16S rRNA gene amplicon sequencing as previously described (Zhang et al. 2024). DNA were from 0.5g of each soil using the Fast DNA®SPIN Kit for Soil (Q-BIOgene, Carlsbad, CA, USA) according to the manufacturer’s instructions. The DNA extract was checked on 1% agarose gel, and DNA concentration were determined with Invitrogen Qubit 4.0 (Thermo Fisher Scientific, America) Qubit^TM^ dsDNA HS Assay Kit (Thermo Fisher Scientific, America).

The hypervariable regions V4−V5 of the archaeal 16S rRNA gene were amplified with primer sets (Arch344F: 5’-ACGGGGYGCAGCAGGCGCGA-3’, Arch934R: 5’-GTGCTCCCCCGCCAATTCCT-3’) (Casamayor et al. 2002) by using an ABIGeneAmp®9700 PCR thermocycler (ABI, CA, USA). The PCR amplification was performed as follows: initial denaturation at 95°C for 3 min, followed by 27-30 cycles of denaturing at 95[ for 30 s, annealing at 55°C for 30 s, extension at 72°C for 45 s, single extension at 72°C for 10 mins, and ending at 10°C. The PCR mixtures contained: 4 μL 5 × *TransStart*Fastpfu buffer, 2 μL 2.5 mM dNQTPs, 0.8 μL 5 μM forward primer, 0.8 μL 5 μM reverse primer, 0.4 μL *TransStart*Fastpfu DNA Polymerase, 10 ng template DNA, with ddH_2_O added to increase the volume to 20 μL. The PCR procedures were performed in triplicate. The PCR product was extracted from 2% agarose gel and purified using the AxyPrep DNA Gel Extraction Kit (Axygen Biosciences, Union City, CA, USA) in accordance with the manufacturer’s instructions and quantified using a Quantus^TM^ Fluorometer (Promega, USA). Purified amplicons were pooled in equimolar and paired-end sequences on an Illumina NovaSeq PE250 platform (Illumina, San Diego, USA).

The sequencing reads were processed using the USEARCH pipeline(Schloss Patrick et al. 2009). Paired-end reads were merged, and then the primers were removed. Sequences were quality-screened with the following settings: Those with a maximum expected error probability of >1.0 were removed from the analysis. Thereafter, all sequences were classified into zero-radius operational taxonomic units (zOTUs) at 100% nucleotide sequence identity(Schloss Patrick et al. 2009). The sequences were classified using Wang’s method against the Silva database (2019.12, release 138), with a minimum confidence score of 80%(Wang et al. 2007). Analysis of sequencing data was based on zOTUs. Sequence data associated with this project have been deposited in the NCBI Short Read Archive Database with Accession Number: PRJNA1173973.

### Quantitative polymerase chain reaction (qPCR) assays

The copy numbers of bacterial 16S rRNA gene, archaeal 16S rRNA gene and *mcrA* gene coding for the alpha subunit of the methyl CoM reductase were quantified using qPCR with primer sets of 338F/518R(Bao et al. 2018) (338F: 5’-ACTCCTACGGGAGGCAGCAG-3’, 518R: 5’-ATTACCGCGGCTGCTGG-3’), 364F/934R (364F: 5’-CGGGGYGCASCAGGCGCGAA-3’, 934R: 5’-GCGCTCCCCCGCCAATTCCT-3’) (Casamayor et al. 2002) and maIs-mod-F/mcrA-rev-R(Zhang et al. 2020a) (maIs-mod-F: 5’-GGYGGTGTMGGDTTCACMCARTA-3’, mcrA-rev-R: 5’-CGTTCATBGCGTAGTTVGGRTAGT-3’), respectively. The construction of qPCR standards was carried out as previously described(Liu et al. 2018). Quantitative PCR amplification was performed on a Thermofisher ABI QuantStudio 1 (Thermo Fisher Scientific, America). The 20 μL reaction mixture contained 2 μL of the DNA template, 10 μL of TB GreenTM Premix EX TaqTM[ (TaKaRa), 0.5 μL of each forward and reverse primer (20 µM), 0.4 μL ROX Reference Dye (TaKaRa), and 6.6 μL ddH_2_O. qPCR conditions for 16S rRNA and *mcrA* gene were as follows: initial denaturation (94°C, 5 min), followed by 40 cycles of denaturation (94°C, 30 s), annealing (52°C, 30 s) and elongation and data collection (72°C, 1 min). A melting curve analysis was conducted to confirm the specificity of the PCR products. Each measurement was performed in triplicate.

### Calculations of methane and CO_2_ production rate and the temperature coefficient (Q10)

The Q10 for CH_4_ and CO_2_ production were calculated as follows(Duc et al. 2010):

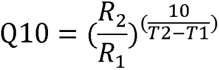

T1 and T2 are the incubation temperature used, here T1=5°C and T2=15°C. R1 and R2 are average rates of CH_4_ or CO_2_ production at T1 and T2, respectively. The average rate of CH_4_ or CO_2_ production is calculated as follows: During the incubation, the approximate linear increase phase of methane and CO_2_ concentrations was used to calculate the average rate of methane and CO_2_ production according to the accumulation of methane and CO_2_ in each incubation tube. The average production rate of methane and CO_2_ was obtained by establishing a linear correlation between methane concentration or CO_2_ and time during this period (including at least 3 measurement points) and calculating the slope of the fitting line. Since the methane production rate of different culture bottles at some sites was too low to calculate the production rate at 5°C, its Q10 could not be calculated (see the results section for more details).

### Statistical analysis and figures generation

All statistical analyses were performed using R version 3.9.2 (R Development Core Team, 2019). Alpha diversity metrics were calculated using the “picante” package and “vegan” package in the software R. The Mantel test was carried out in R with the mantel function in the package “ggcor” to evaluate the relationships between methane and carbon dioxide production rates, Q10, and influencing variables. The order of feature importance was determined using a random forest (RF) in the R package “randomForest” over 1000 iterations(Zhang et al. 2018). dbRDA ordination analysis of the archaeal community structures in sediments, based on Bray-Curtis distance, was conducted to identify the differences in these community structures before and after incubation. The analysis of variance was performed to test significant differences between treatments using the DUNCAN test within “agricolae” package in R.

## Results

### Characteristics of the GFS sediments

Sediment characteristics showed considerable variations and spatial heterogeneity. The concentrations of carbon (C), nitrogen (N), and phosphorus (P) were extremely low. Specifically, the levels of TOC, TN, NH_4_^+^-N, NO_3_^-^-N and TP ranged from 475.0 to 4,084.1 μg/g dry weight sediments (DWS), 0 to 826.9 μg/g DWS, 2.6 to 31.3 μg/g DWS, 0 to 1.3 μg/g DWS and 6.4 to 236.4 μg/g DWS, respectively (Fig. 1b). The C to N ratio (C/N) varied between 4.7 and 19.1. The sediments were slightly alkaline, with pH values ranging from 7.89 to 9.27 (Fig. 1b), and EC ranging from 36.3 to 178.0 μs/cm. Most factors showed an obvious negatives correlation with altitude, although not significant (Fig. S2), while only pH (*P*=0.01) was significantly positively correlated with altitude (Fig. S2).

### Rates and Q10 of gas production

Substantial accumulation of methane (Fig. 2) and CO_2_ (Fig. S3) were observed for all samples (except for site R7, 5,145 m) incubated at both temperatures. The total amount of CH_4_ and CO_2_ accumulated in the headspace were in a range of 0.01-5.71 and 0.33-1.74 µmol, respectively (Fig. 2; Fig. S3). The ratio of accumulated CH_4_ to CO_2_ were 0.05-2.76 (median) and 0.76-3.91 (median) at 5°C and 15°C (Fig. S4), respectively. Total carbon release in the form of CH_4_ and CO_2_ accounted for 0.0002%-0.04% and 0.003%-0.01% of sediments TOC at 5°C and 15°C, respectively (Fig. S4).

**Fig. 2.**
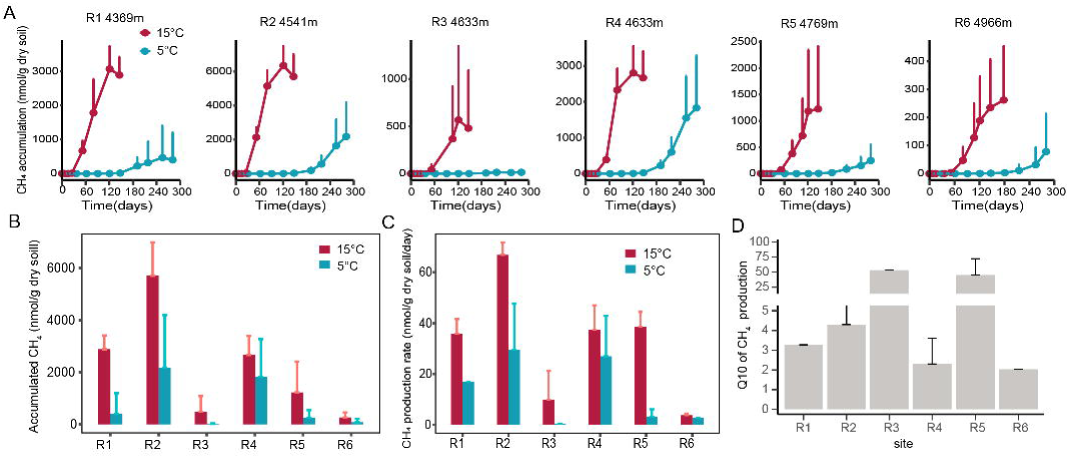
Accumulation of CH4, rate and Q10 of methane production. a. dynamics of methane accumulation in the headspace of incubation tubes. b. Total amount of methane accumulated in the headspace. Data shown are means ± SD, n=2-4 for a and b. c. Rate of CH_4_ production. d. Q10 of methane production.

The rate of CH_4_ production were 0.34-29.59 and 3.80-66.83 nmol/g dry soil day^-1^ at 5°C and 15°C, respectively (Fig. 2). And the rate of CO_2_ production were 1.86-7.52 and 2.0-24.10 nmol/g dry soil day^-1^ at 5°C and 15°C, respectively (Fig. S3). For most sites, the amount of accumulated methane and CO_2_ (Fig. 2 and Fig. S3) and rate of methane and CO_2_ production (Fig. 2 and Fig. S3) were significantly higher at 15°C than those at 5°C.

With the average rate of methane and CO_2_ production at 5°C and 15°C, we calculated the temperature coefficient (Q10) for both processes (Fig. 2 and Fig. S3). Q10 were between 1.22 and 57.15 and between 1.57 and 13.15 for methane and CO_2_ production, respectively. In addition, at 15°C but not at 5°C, we observed significant negative correlations between altitude and the production rate and accumulated amount of CH_4_ and CO_2_, respectively (Fig. S5). While only the Q10 of CO_2_ production but not CH_4_ showed a significant negative correlation with altitude (Fig. S5).

### Abundance of microbial communities

The copy number of bacterial and archaeal 16S rRNA genes, as well as *mcrA* gene for methanogens, ranged between 3.1×10^7^ and 5.9×10^9^, 9.7×10^4^ and 1.8×10^7^ and 1.8×10^5^ and 2.4×10^6^ copies/g DWS, respectively, in the original samples (Fig. 3). Following incubation, the copy numbers of all three genes increased significantly (*P*<0.001; Fig. 3) in samples across all sites (except for R7). The copy number in post-incubation samples reached 9.8×10^7^-3.5×10^9^, 2.1×10^7^-8.2×10^8^ and 2.1×10^7^-1.2×10^9^ for bacterial 16S rRNA gene, archaeal 16S rRNA gene and *mcrA* gene, respectively. The copies of bacterial 16S rRNA gene increased by more than 10-fold for most sites (Fig. 3a), while the archaeal 16S rRNA gene and *mcrA* gene increased by more than 10- to 1,000-folds (Fig. 3b and c). Copies of all three genes were similar between samples incubated at 15°C and 5°C, which might be due to much longer incubation at 5°C than those at 15°C before sampling. Besides, only a slightly higher number of gene copies were observed in samples from lower altitudes (Fig. 3).

**Fig. 3.**
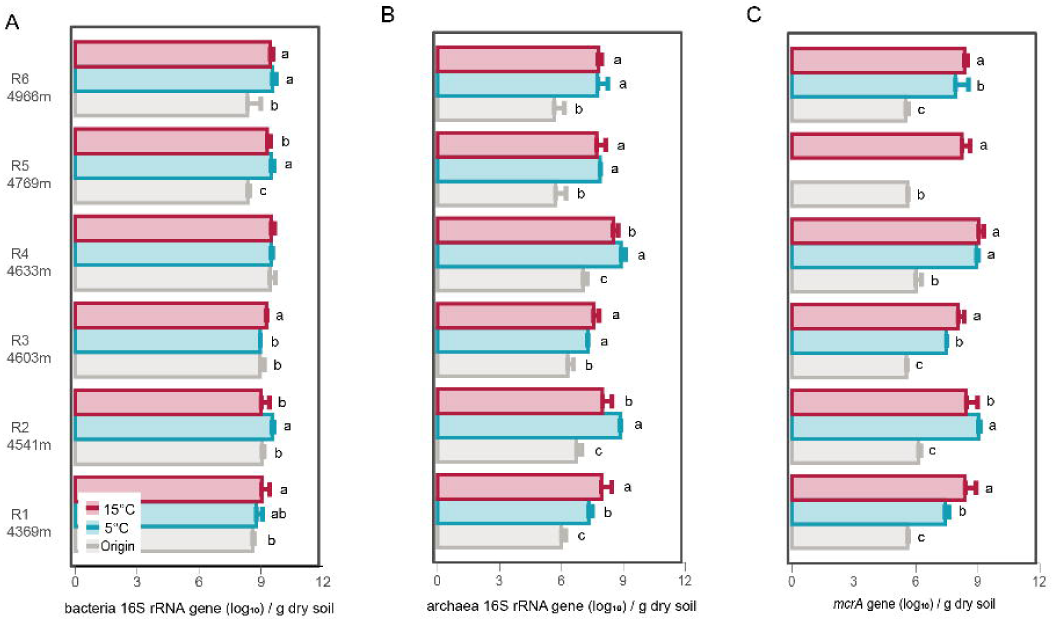
Copy numbers of bacterial and archaeal 16S rRNA genes and *mcrA* gene in samples before and after incubation. a. bacteria 16S rRNA gene, b. archaea 16S rRNA gene and c. *mcrA* gene.

### Community structures of Archaea

In the original samples, *Nitrosophaeraceae* dominated the archaeal community (median relative abundance of 98.64%) (Fig. 4). Methanogenic taxa, including *Methanosarciniaceae*, *Methanotrichaceae* (formerly *Methanosaeceae*), *Methanobacteriaceae,* and *Methanomicrobiales* (including *Methanoregulaceae*, *Methanospirillaceae*, and unclassified *Methanomicrobiales*) represented a minor fraction, with relative abundances of less than 1% across all samples. *Metanomassiliicoccaceae* were detected at only one site (R1), and *Methanocellaceae* were not detected in the original samples (Fig. 4).

**Fig. 4.**
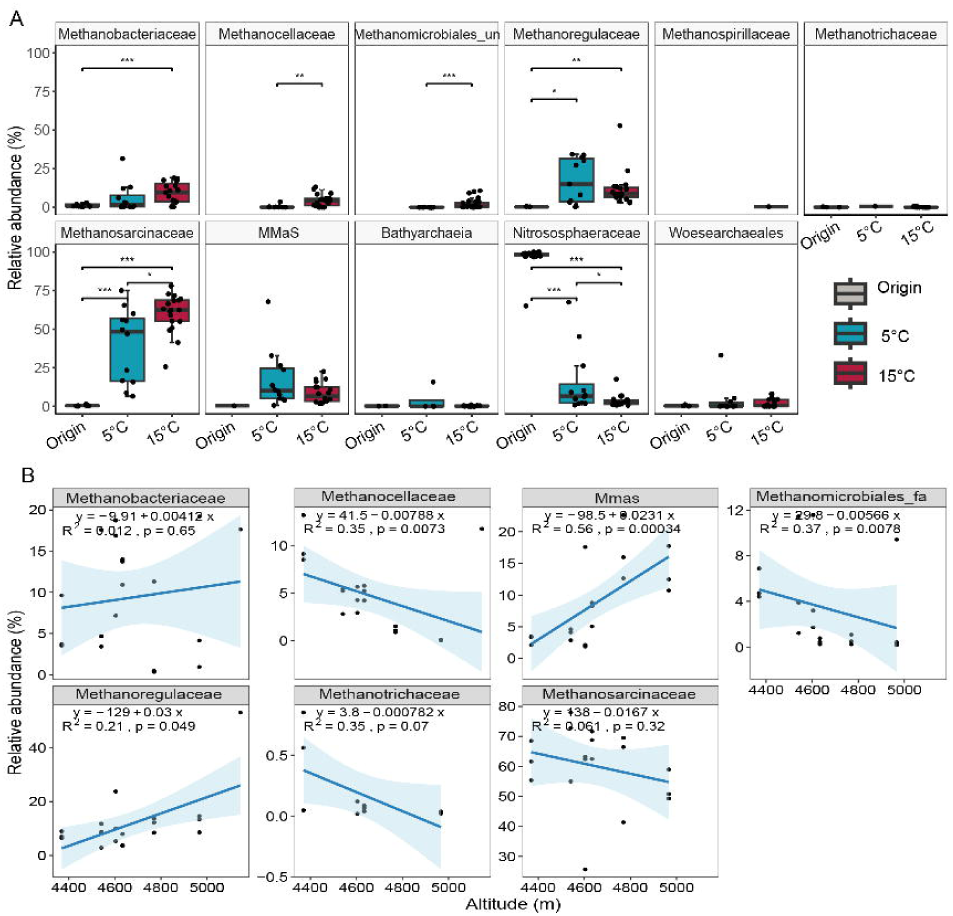
Relative abundance of major archaeal families and their correlations with altitude. a boxplot showing the relative abundance of major archaeal families. b correlation of methanogens family in samples after incubation at 15°C with altitude. See Fig. S6 for the original and post-incubation samples at 5°C. Abbreviation, MMsa *Methanomassiliicoccaceae*.

After 280 and 140 days of incubation at 5°C and 15°C, respectively, the relative abundance of all methanogens increased profoundly (Fig. 4a), especially at 15°C. *Methanosarcinaceae*, the facultative acetoclastic methanogens, become the dominant archaeal groups (with relative abundance >55%) at both temperatures and has a significantly higher relative abundance at 15°C compared to 5°C. In contrast, *Methanotrichaceae*, obligate acetoclastic methanogens, maintained low relative abundance (<1%) and exhibited no significant change after incubation at both temperatures. Hydrogenotrophic methanogens, including *Methanocellaceae*, *Methanobacteriaceae*, *Methanoregulaceae*, *Methanospirillaceae*, and unclassified *Methanomicrobiales*, showed a significant increase (P<0.05) at 15°C compared to the original samples. Notably, only *Methanocellaceae* and unclassified *Methanomicrobiales* exhibited a significant increase (P<0.05) at 15°C compared to 5°C. The relative abundance of *Methanobacteriaceae* was significantly higher at 15°C compared to the original samples (*P*<0.05) but not those at 5°C. *Methanomassiliicoccaceae*, the methylotrophic methanogens, which were 25.4 and 16.5 folds higher in post-incubation samples at 15°C and 5°C than those of the original samples, respectively (Fig. 4a), displayed similar relative abundances at both temperatures. Additionally, the relative abundance of ammonia-oxidizing archaea, *Nitrososphaeraceae*, decreased significantly after incubation at both temperatures, especially at 15°C. Meanwhile, families within *Bathyarchaeia* and *Woesearchaeales* remained relatively unchanged between the original and post-incubation samples (Fig. 4a).

To find if specific taxonomic groups varied with altitude, we conducted a correlation analysis between the relative abundance of methanogenic families and altitude (Fig. 4b and Fig. S6). In both the original samples and post-incubation samples at 5°C, no significant altitude-dependent trends across all families were observed, except for *Methanobacteriaceae* at 5°C (Fig. S6). However, after incubation at 15°C, several groups exhibited significant correlations with altitude. The correlation between hydrogenotrophic methanogens and altitude was variable: *Methanocellaceae* and unclassified *Methanomicrobiales* showed significant negative correlations, while *Methanoregulaceae* displayed a significant positive correlation with altitude (Fig. 4b; all *P*<0.05). The methylotrophic methanogens, *Methanomassiliicoccaceae*, displayed a significant positive correlation with altitude (Fig 4b; *P*<0.05). In contrast, the relative abundance of acetoclastic methanogens, including both *Methanotrichaceae* and *Methanosarcinaceae*, were only slightly decreased with increasing altitude.

### Alpha-diversity, beta-diversity and factors shaping the sediment archaeal communities

The alpha-diversity of sediment archaea was notably low in both the original and post-incubation samples, with median values ranging between 22 and 63 for Richness, 0.36 and 0.46 for Peilou’s Evenness, and 1.92 and 2.33 for Shannon’s H (Fig. 5), respectively. Incubation at 15°C led to a significant increase in both Richness (*P*<0.05) and Shannon’s H (*P*<0.05), but a significant decrease in Evenness (*P*<0.05). Similarly, incubation at 5°C significantly increased Richness (*P*<0.05) and caused a significant decrease in Evenness (*P*<0.05), but with no significant effects on Shannon’s H (Fig. 5). Key factors influencing alpha-diversity indices included altitude, pH, TN, TOC, C/N ratio (Fig. S7). Notably, altitude was significantly negatively correlated with both Shannon and Richness indices in the original and 15°C post-incubation samples. TN and TOC exhibited significant positive correlations with Richness indices across both original and post-incubated samples at both temperatures, except for TOC at 15°C. Additionally, TN showed a significant positive correlation with the Shannon indices of post-incubated samples at 5°C (Fig. S7).

**Fig. 5.**
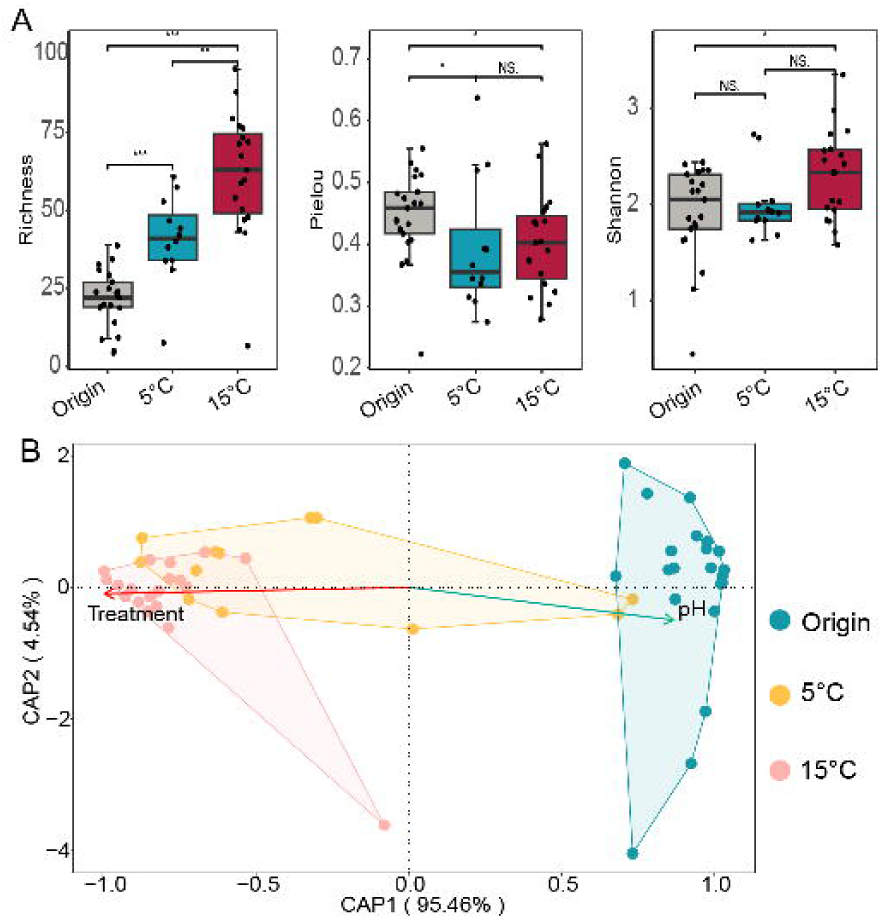
Diversity of archaeal communities. a. alpha-diversity of archaeal communities in the original, post-incubation samples at 5°C and 15°C, respectively. b. betadiversity.

The overall composition of the archaeal community was significantly impacted by incubation (ANOSIM, *P*<0.001) (Fig. 5b). In the dbRDA ordination plot, post-incubation samples at 5°C and 15°C, along with the original samples, formed distinct clusters, although some overlap was observed between the 5°C and 15°C treatments. In addition to temperature, pH also played a crucial role in shaping archaeal communities, as incubation at both 5°C and 15°C significantly decreased slurry pH (Fig. S8). We further investigated the factors influencing archaeal communities beyond incubation temperatures. In both the original and post-incubation samples at 15°C, altitude and pH emerged as the key factors significantly correlated with archaeal communities, as determined by the Mantel test (Fig. 5 and Fig. S9). Conversely, at 5°C, only altitude showed significant correlations with archaeal communities (Fig. 5 and Fig. S9).

### Factors influence gas production and Q10

For CH_4_ production rates, significant correlations with the rate of CH_4_ production at 5°C were observed for pH, EC and C/N (Fig. 6a). At 15°C, significant factors included altitude, pH and C/N (Fig. 6b). Regarding CO_2_ production at 5°C, pH, EC, and TP showed significant correlations with the rate (Fig. S10a). At 15°C, altitude, pH, EC, TP, and C/N, were identified as significant factors (Fig. S10b). For Q10, altitude, pH and C/N were significantly correlated with the Q10 of CO_2_ production (Fig. S10c). In contrast, no factors were significantly correlated with the Q10 of CH_4_ production (Fig. 6c). For the total amount of CH_4_ accumulated in the headspace, significant correlations were observed with pH, EC, and C/N at 5°C, and with altitude, pH and C/N at 15°C. For the total amount of accumulated CO_2_, significant factors included pH, EC, and TP at 5°C, and altitude, pH, EC, TP, and C/N at 15°C (Fig. S11).

**Fig. 6.**
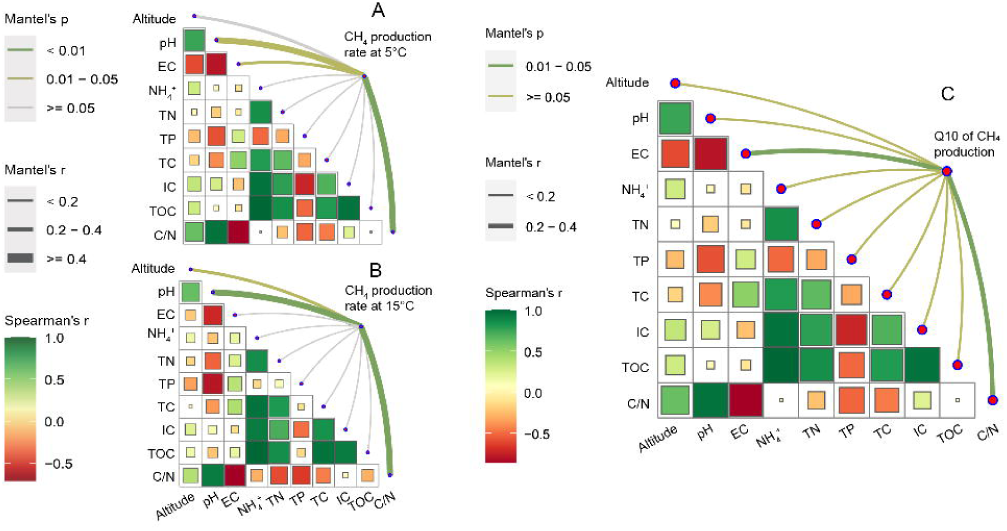
Factors influence methane production rate and Q10. Mantel test analysis of sediment properties with methane production rate at 5 °C (a), and 15 °C (b) and Q10 of methane production (c), respectively.

## Discussion

Here, we conducted the first study on the GFS methanogenic archaeal community, methane production, and their responses to temperature. Through deep amplicon sequencing, we found all major types of methanogenic archaea, including hydrogenotrophic, acetoclastic, and methylotrophic groups. These groups are well-documented in other high-altitude, non-glacier-fed anaerobic ecosystems on the Tibetan Plateau, including lake sediments(Liu et al. 2016b), riverbed sediments on the east Tibetan plateau(Zhang et al. 2020b), thermokarst lake sediments(Yang et al. 2023), and wetlands(Xing et al. 2022, Yang et al. 2017). Hence, methanogens living in GFS are as diverse as other non-glacier-fed anaerobic ecosystems of the Tibetan Plateau. However, in the original sample, the abundance of all methanogenic archaea (10^4^∼10^5^) and relative abundance are very low (all <2%). These values align with reports from headwater river sediments and glacier meltwater sediments in the Tibetan Plateau(Du et al. 2024, Zhang et al. 2020b). Given that our sampling occurred in May (Fig. S1) when river temperatures were low (4°C–8°C), and that high turbulence likely oxygenated the sediments (Fig. S1), we hypothesized that methanogenic archaea were dormant, resulting in low abundance. This is further supported by the long lag phase observed before obvious methane production occurred during incubation at both temperatures (Fig. 2). Outside the Tibetan Plateau, Michoud et al. detected only *Methanomicrobiales* and *Methanotrichales* in the Mt. Stanley GFS in Africa. We propose that a similar diversity of methanogens exists in Mt. Stanley GFS as Rongbu GFS, but metagenomic binning, which captures high-abundance genomes, may have limited their detection(Michoud et al. 2023).

Our results showed that under elevated temperatures, methanogenic archaea in GFS have very high methane production potential, which was comparable to most previously studied river sediments(Mach et al. 2015). Alongside methane production, the copy numbers of the archaeal 16S rRNA gene and *mcrA* gene increased by 10^3^-10^4^ times, indicating that these microorganisms are capable of rapid proliferation under favorable conditions. This data indirectly supports that sediment methanogenesis is a key contributor to methane emissions observed in GFS in glacier melting seasons(Du et al. 2024, Zhang et al. 2021b), with the methanogenesis process likely occurring in micro-anaerobic “pocket”, where methane accumulates and is released as bubbles through ebullition. This supports previous findings showing ebullition as the dominant methane emission pathway in headwater streams on the Tibetan Plateau, accounting for up to 79% of total methane flux(Zhang et al. 2020b).

As temperature rises, facultative acetoclastic methanogens, particularly *Methanosarcinaceae*, exhibit the most significant increase in relative abundance (Fig. 4). This shift suggests that acetoclastic methanogenesis may become the dominant pathway for methane production in GFS at elevated temperatures. This is consistent with Du’s isotope observations in glacier meltwater sediments(Du et al. 2024): the carbon isotopic compositions of CH_4_ and CO_2_ in the ice cave and terminus meltwater indicated δ^13^C-CH_4_ depletion compared to ambient air, suggesting an acetate fermentation pathway(Du et al. 2024). Moreover, this is also aligned with the findings of Arctic subglacial ecosystems, of which acetoclastic methanogensis dominates the methane production pathways (Lamarche-Gagnon et al. 2019a). Therefore, these findings highlight the importance of acetoclastic methanogenesis in glacier subglacial and proglacial ecosystems. In addition, the methanogenic archaeal community in GFS exhibited a significant altitude dependence with increasing temperatures (Fig. 4b). Specifically, methylotrophic methanogens, *Methanomassiliicoccaceae*, showed a positive correlation with altitude (*P*<0.001; Fig. 4b), reaching up to 15-20% of the archaeal community at elevations above 4,600 m. This may be due to the physiological adaptations of *Methanomassiliicoccaceae*, including trehalose biosynthesis for temperature regulation, agmatine production for pH regulation, and overall tolerance to low temperatures, high pH, and oxygen exposure(Borrel et al. 2014, Chen et al. 2023, Söllinger and Urich 2019). Therefore, we propose that in the future, as temperature rises, methylotrophic methanogenesis will play a key role in the high-altitude GFS of Mt. Everest.

Although we found significant influences of incubation temperature and altitude on archaeal community composition, TOC and TN did not show obvious effects, which contrasts with what has been observed in previous studies of river sediments(Bednařík et al. 2019, Comer-Warner et al. 2018). It may be related to the fact that GFS is an extremely oligotrophic environment(Kohler et al. 2024), the concentration of TOC and TN in all original samples (Fig. 1) falls far below the requirement of methanogens. Therefore, methanogens in GFS do not show responses to the variations of these environmental factors. However, preliminary experiments (unpublished data from our group) indicate that adding substantial substrates (0.5% of cellulose) can significantly increase methane production and methanogen abundance, suggesting TOC can influence methanogens under more nutrient-rich conditions, which is consistent with previous studies on river sediment methanogenesis(Bodmer et al. 2020, Romeijn et al. 2019). Kohler et al. (2024)(Kohler et al. 2024) predict an increase in GFS primary productivity due to a “green transition” towards autotrophy in alpine ecosystems, which could increase organic matter in stream sediments, potentially making organic matter a key regulator of future methane emissions in GFS.

Finally, we revealed factors influencing the rate and temperature sensitivity of the GFS carbon release process under anaerobic conditions. Similar to changes in methanogen communities, methane and CO_2_ production rates were negatively correlated with altitude, with higher altitudes exhibiting lower rates. Therefore, as temperature rises, methane and CO_2_ emissions from low-altitude areas will likely increase more than from high-altitude areas, contributing to spatial heterogeneity in carbon emissions. The temperature sensitivity of methane and CO_2_ production is similar to the reported range of CH_4_ and CO_2_ production from river sediments (Comer-Warner et al. 2018, Rodriguez et al. 2018, Wilkinson et al. 2019) and showed great variations. The temperature sensitivity of CO_2_ production also showed a negative correlation with altitude, while that of methane production did not. Consequently, CO_2_ emissions from low-altitude GFS may increase more rapidly than those from high-altitude regions, further amplifying the spatial variability of carbon release in alpine GFS basins.

In conclusion, this study advances our understanding of methanogenic archaea in glacial-fed streams. The Rongbuk GFS sediments harbor a diverse and abundant community of methanogenic archaea, including hydrogenotrophic, acetoclastic, and methylotrophic types. These microorganisms can proliferate and produce substantial amounts of methane under rising temperatures, indicating that GFS may play a critical role in methane cycling within glacial basins. Our results underscore the need to consider greenhouse gas emissions from alpine GFS in regional and global methane and carbon dioxide budgets, especially as the climate continues to warm.

## Supporting information

Supplementary Materialss

## Acknowledgements

We thank Yubo Su and Shuai Yuan for their help on field sampling. This work was supported by the National Natural Science Foundation of China General Program (42171144), the National Natural Science Foundation of China for Excellent Young Scientists Fund Program (42222105), the Second Tibetan Plateau Scientific Expedition and Research Program (STEP) (2021QZKK0100).

## Data Availability

Amplicon sequencing data of Archaeal 16S rRNA genes from the current study were deposited at NCBI with accession number of PRJNA1173973.

## Author contributions

Pengfei Liu initiated and designed the study; Weizhen Zhang, Peihua Shen, and Jingyi Zhu carried out sample processes, and Peihua Shen carried out sediment microcosm incubation experiment, CH_4_, CO_2_ and pH measurement, DNA extraction; Pengfei Liu, Wei Ma, Miao Lin, Shaoyi Tian, Peihua Shen and Hongfei Chi performed all data curation, analysis and visualization, with results interpretations from also Junzhi Liu. All authors contributed to the editing of the text and approved the final version.

## Reference

Acuna V, Wolf A, Uehlinger U et al. Temperature dependence of stream benthic respiration in an Alpine river network under global warming. Freshwater Biology 2008;53: 2076–88.

Angle JC, Morin TH, Solden LM et al. Methanogenesis in oxygenated soils is a substantial fraction of wetland methane emissions. Nat Commun 2017;8: 1567.

Bao Y, Li B, Xie S et al. Vertical profiles of microbial communities in perfluoroalkyl substance-contaminated soils. Annals of Microbiology 2018;68: 399–408.

Battin TJ, Besemer K, Bengtsson MM et al. The ecology and biogeochemistry of stream biofilms. Nat Rev Microbiol 2016;14: 251–63.

Battin TJ, Lauerwald R, Bernhardt ES et al. River ecosystem metabolism and carbon biogeochemistry in a changing world. Nature 2023;613: 449–59.

Bednarik A, Blaser M, Matousu A et al. Effect of weir impoundments on methane dynamics in a river. Sci Total Environ 2017;584: 164–74.

Bednařík A, Blaser M, Matoušů A et al. Sediment methane dynamics along the Elbe River. Limnologica 2019;79: 125716.

Bodmer P, Wilkinson J, Lorke A. Sediment Properties Drive Spatial Variability of Potential Methane Production and Oxidation in Small Streams. J Geophys Res Biogeosci 2020;125: e2019JG005213.

Borrel G, Parisot N, Harris HMB et al. Comparative genomics highlights the unique biology of Methanomassiliicoccales, a Thermoplasmatales-related seventh order of methanogenic archaea that encodes pyrrolysine. BMC Genomics 2014;15: 679.

Brablcová L, Buriánková I, Badurová P et al. Methanogenic archaea diversity in hyporheic sediments of a small lowland stream. Anaerobe 2015;32: 24–31.

Busi SB, Bourquin M, Fodelianakis S et al. Genomic and metabolic adaptations of biofilms to ecological windows of opportunity in glacier-fed streams. Nat Commun 2022;13: 2168.

Casamayor EO, Massana R, Benlloch S et al. Changes in archaeal, bacterial and eukaryal assemblages along a salinity gradient by comparison of genetic fingerprinting methods in a multipond solar saltern. Environ Microbiol 2002;4: 338–48.

Chaudhary PP, Rulík M, Blaser M. Is the methanogenic community reflecting the methane emissions of river sediments?—comparison of two study sites. MicrobiologyOpen 2017;6: e00454.

Chen X, Xue D, Wang Y et al. Variations in the archaeal community and associated methanogenesis in peat profiles of three typical peatland types in China. Environmental Microbiome 2023;18: 48.

Comer-Warner SA, Romeijn P, Gooddy DC et al. Thermal sensitivity of CO2 and CH4 emissions varies with streambed sediment properties. Nat Commun 2018;9.

Conrad R. The global methane cycle: recent advances in understanding the microbial processes involved. EnvironMicrobiol Rep 2009;1:285–92.

Conrad R. Importance of hydrogenotrophic, aceticlastic and methylotrophic methanogenesis for methane production in terrestrial, aquatic and other anoxic environments: A mini review. Pedosphere 2020;30: 25–39.

Conrad R. Complexity of temperature dependence in methanogenic microbial environments. Front Microbiol 2023;14.

Conrad R, Ji Y, Noll M et al. Response of the methanogenic microbial communities in Amazonian oxbow lake sediments to desiccation stress. Environ Microbiol 2014;16: 1682–94.

Crawford JT, Loken LC, West WE et al. Spatial heterogeneity of within-stream methane concentrations. Journal of Geophysical Research-Biogeosciences 2017;122: 1036–48.

Du Z, Cui H, Wang L et al. Characteristics of methane and carbon dioxide in ice caves at a high-mountain glacier of China. Sci Total Environ 2024;946: 174074.

Duc NT, Crill P, Bastviken D. Implications of temperature and sediment characteristics on methane formation and oxidation in lake sediments. Biogeochemistry 2010;100: 185–96.

Gao T, Zhang Y, Wei D et al. Cryospheric melting enhances methane emissions from inland waters on the Tibetan Plateau. 2023.

King O, Bhattacharya A, Ghuffar S et al. Six Decades of Glacier Mass Changes around Mt. Everest Are Revealed by Historical and Contemporary Images. One Earth 2020;3: 608–20.

Kohler TJ, Bourquin M, Peter H et al. Global emergent responses of stream microbial metabolism to glacier shrinkage. Nat Geosci 2024, DOI 10.1038/s41561-024-01393-6.

Konya K, Sueyoshi T, Iwahana G et al. CH4 emissions from runoff water of Alaskan mountain glaciers. Sci Rep 2024;14: 10558.

Lamarche-Gagnon G, Wadham JL, Sherwood Lollar B et al. Greenland melt drives continuous export of methane from the ice-sheet bed. Nature 2019a;565: 73–7.

Lamarche-Gagnon G, Wadham JL, Sherwood Lollar B et al. Greenland melt drives continuous export of methane from the ice-sheet bed. Nature 2019b;565: 73–7.

Li M, Peng C, Zhang K et al. Headwater stream ecosystem: an important source of greenhouse gases to the atmosphere. Water Research 2021;190: 116738.

Li M, Shi G, Li Y et al. Isotopic Constraints on Sources and Transformations of Nitrate in the Mount Everest Proglacial Water. Environ Sci Technol 2023;57: 20844–53.

Liu P, Pommerenke B, Conrad R. Identification of *Syntrophobacteraceae* as major acetate-degrading sulfate reducing bacteria in Italian paddy soil. Environ Microbiol 2018;20: 337–54.

Liu Y, Priscu JC, Xiong J et al. Salinity drives archaeal distribution patterns in high altitude lake sediments on the Tibetan Plateau. Fems Microbiology Ecology 2016a;92.

Liu Y, Vick-Majors TJ, Priscu JC et al. Biogeography of cryoconite bacterial communities on glaciers of the Tibetan Plateau. FEMS Microbiology Ecology 2017a;93.

Liu Y, Whitman WB. Metabolic, Phylogenetic, and Ecological Diversity of the Methanogenic Archaea. Ann N Y Acad Sci 2008;1125: 171–89.

Liu Y, Yao T, Gleixner G et al. Methanogenic pathways, 13C isotope fractionation, and archaeal community composition in lake sediments and wetland soils on the Tibetan Plateau. J Geophys Res Biogeosci 2013;118: 650–64.

Liu YQ, Conrad R, Yao TD et al. Change of methane production pathway with sediment depth in a lake on the Tibetan plateau. Palaeogeography Palaeoclimatology Palaeoecology 2017b;474: 279–86.

Liu YQ, Priscu JC, Xiong JB et al. Salinity drives archaeal distribution patterns in high altitude lake sediments on the Tibetan Plateau. Fems Microbiology Ecology 2016b;92.

Mach V, Blaser MB, Claus P et al. Methane production potentials, pathways, and communities of methanogens in vertical sediment profiles of river Sitka. Front Microbiol 2015;6.

Michoud G, Kohler TJ, Ezzat L et al. The dark side of the moon: first insights into the microbiome structure and function of one of the last glacier-fed streams in Africa. Royal Society Open Science 2023;10: 230329.

Milner AM, Khamis K, Battin TJ et al. Glacier shrinkage driving global changes in downstream systems. Proc Natl Acad Sci U S A 2017;114: 9770–8.

Nagler M, Praeg N, Niedrist GH et al. Abundance and biogeography of methanogenic and methanotrophic microorganisms across European streams. Journal of Biogeography 2021;48: 947–60.

Rocher-Ros G, Stanley EH, Loken LC et al. Global methane emissions from rivers and streams. Nature 2023, DOI 10.1038/s41586-023-06344-6.

Rodriguez M, Gonsiorczyk T, Casper P. Methane production increases with warming and carbon additions to incubated sediments from a semiarid reservoir. Inland Waters 2018;8: 109–21.

Romeijn P, Comer-Warner SA, Ullah S et al. Streambed Organic Matter Controls on Carbon Dioxide and Methane Emissions from Streams. Environ Sci Technol 2019;53: 2364–74.

Saunois M, Stavert AR, Poulter B et al. The Global Methane Budget 2000–2017. Earth Syst Sci Data 2020;12: 1561–623.

Schloss Patrick D, Westcott Sarah L, Ryabin T et al. Introducing mothur: Open-Source, Platform-Independent, Community-Supported Software for Describing and Comparing Microbial Communities. Appl Environ Microb 2009;75: 7537–41.

Söllinger A, Urich T. Methylotrophic methanogens everywhere — physiology and ecology of novel players in global methane cycling. Biochemical Society Transactions 2019;47: 1895–907.

Song C, Liu S, Wang G et al. Inland water greenhouse gas emissions offset the terrestrial carbon sink in the northern cryosphere. Sci Adv 2024;10: eadp0024.

Stanley EH, Casson NJ, Christel ST et al. The ecology of methane in streams and rivers: patterns, controls, and global significance. Ecological Monographs 2016;86: 146–71.

Sun X, Zhang Q, Zhang G et al. Melting Himalayas and mercury export: Results of continuous observations from the Rongbuk Glacier on Mt. Everest and future insights. Water Research 2022;218: 118474.

Wang L, Oien A. Determination of Kjeldahl Nitrogen and Exchangeable Ammonium in Soil by the Indophenol Method. Acta Agr Scand 1986;36: 60–70.

Wang Q, Garrity George M, Tiedje James M et al. Naïve Bayesian Classifier for Rapid Assignment of rRNA Sequences into the New Bacterial Taxonomy. Appl Environ Microb 2007;73: 5261–7.

Wilhelm L, Besemer K, Fragner L et al. Altitudinal patterns of diversity and functional traits of metabolically active microorganisms in stream biofilms. ISME J 2015;9: 2454–64.

Wilhelm L, Singer GA, Fasching C et al. Microbial biodiversity in glacier-fed streams. ISME J 2013;7: 1651–60.

Wilkinson J, Bodmer P, Lorke A. Methane dynamics and thermal response in impoundments of the Rhine River, Germany. Sci Total Environ 2019;659: 1045–57.

Wilkinson J, Maeck A, Alshboul Z et al. Continuous Seasonal River Ebullition Measurements Linked to Sediment Methane Formation. Environ Sci Technol 2015;49: 13121–9.

Xing T, Liu P, Ji M et al. Sink or Source: Alternative Roles of Glacier Foreland Meadow Soils in Methane Emission Is Regulated by Glacier Melting on the Tibetan Plateau. Front Microbiol 2022;13.

Yang G, Zheng Z, Abbott BW et al. Characteristics of methane emissions from alpine thermokarst lakes on the Tibetan Plateau. Nat Commun 2023;14: 3121.

Yang S, Liebner S, Winkel M et al. In-depth analysis of core methanogenic communities from high elevation permafrost-affected wetlands. Soil Biol Biochem 2017;111: 66–77.

Ye Q, Zhang X, Wang Y et al. Monitoring glacier thinning rate in Rongbuk Catchment on the northern slope of Mt. Qomolangma from 1974 to 2021. Ecological Indicators 2022;144: 109418.

Yvon-Durocher G, Allen AP, Bastviken D et al. Methane fluxes show consistent temperature dependence across microbial to ecosystem scales. Nature 2014;507: 488–91.

Zhang C-J, Pan J, Liu Y et al. Genomic and transcriptomic insights into methanogenesis potential of novel methanogens from mangrove sediments. Microbiome 2020a;8: 94.

Zhang J, Zhang N, Liu Y-X et al. Root microbiota shift in rice correlates with resident time in the field and developmental stage. Science China Life Sciences 2018;61: 613–21.

Zhang L, Xia X, Liu S et al. Significant methane ebullition from alpine permafrost rivers on the East Qinghai–Tibet Plateau. Nat Geosci 2020b;13: 349–54.

Zhang Y, Kang S, Wei D et al. Sink or source? Methane and carbon dioxide emissions from cryoconite holds, subglacial sediments, and proglacial river runoff during intensive glacier melting on the Tibetan Plateau. Fundamental Research 2021a, DOI 10.1016/j.fmre.2021.04.005.

Zhang Y, Kang S, Wei D et al. Sink or source? Methane and carbon dioxide emissions from cryoconite holes, subglacial sediments, and proglacial river runoff during intensive glacier melting on the Tibetan Plateau. Fundamental Research 2021b;1: 232–9.

Zhang Z, Liu Y, Liu K et al. Supraglacial and subglacial ecosystems contribute differently towards proglacial ecosystem communities in Kuoqionggangri Glacier, Tibetan Plateau. Communications Earth & Environment 2024;5: 636.

