## Supplementary Materialss for "Exploring methanogenic archaea and their thermal responses in the glacier-fed stream sediments of Rongbuk River Basin, Mt. Everest"

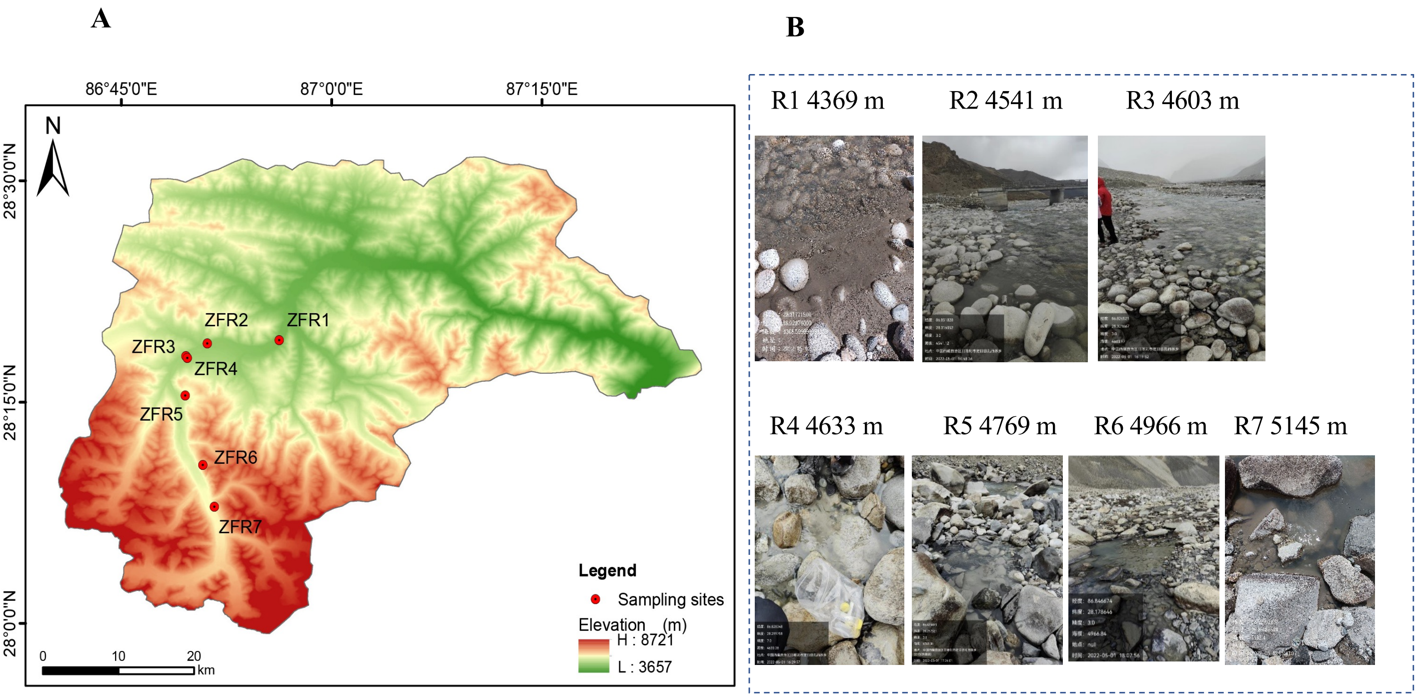


**Fig. S1. Distribution and photos of sampling sites.** (A) Distribution of sampling sites in the River basin. (B) Photos showing the status of glacier-fed stream sediments when sampling.

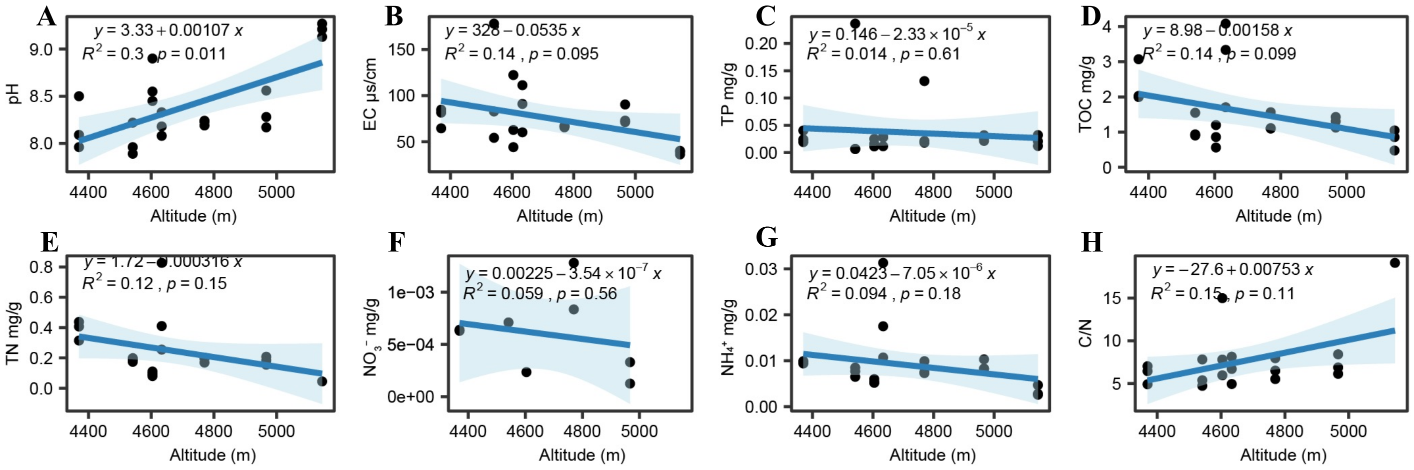


**Fig. S2. Correlation of sediments' physicochemical properties with altitude**. (A) pH. (B) EC. (C) TP, Total phosphor. (D) Total organic carbon, TOC. (E) Total nitrogen, TN. (F) Nitrate, $NO_3^-$. (G) Ammonia, $NH_4^+$. (H) TOC to TN ratio.

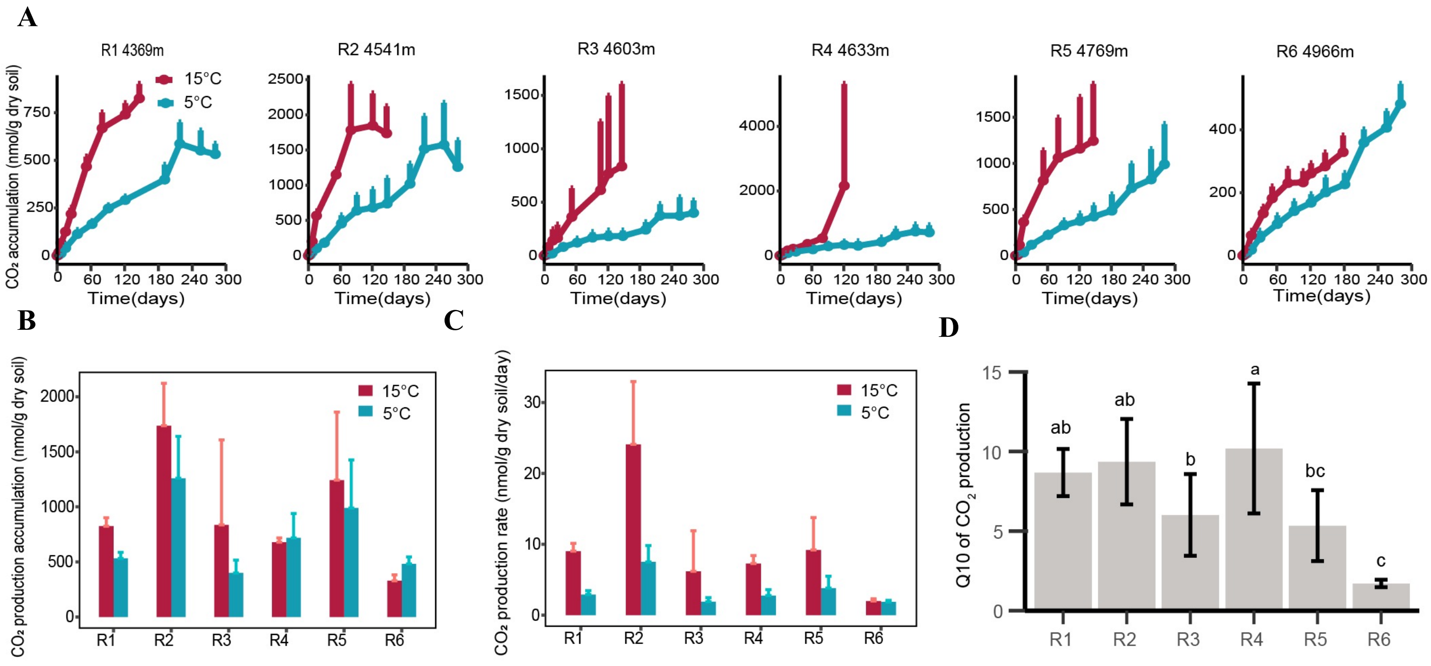


**Fig. S3. Dynamics and Q10 of $CO_2$ production.** (A) Dynamics of $CO_2$ accumulation in the headspace of incubation tubes. (B) The total amount of $CO_2$ accumulated in the headspace at the end of incubation. Data are mean ± SD, n=4 with some exceptions with n=2-3. (C) Rate of $CO_2$ production. (D) Q10 of $CO_2$ production.

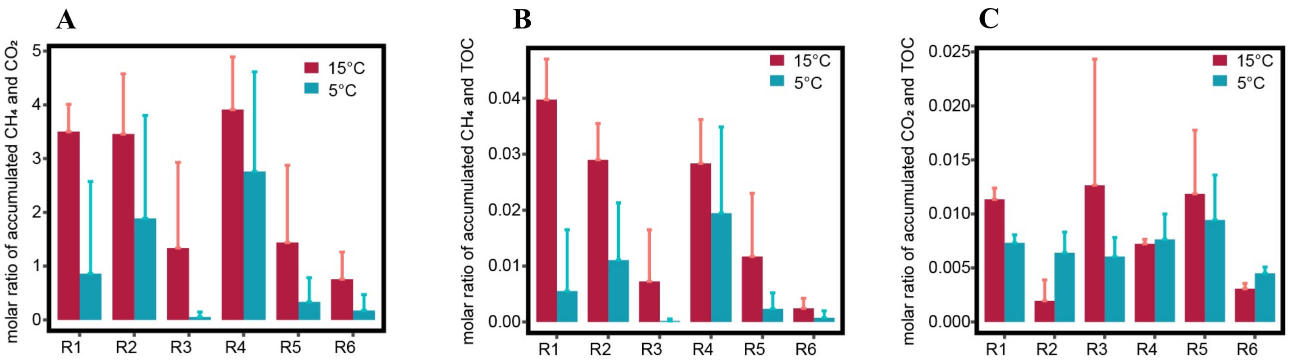


**Fig. S4. Comparison of the released amount of gas and sediment organic carbon.** (A) Molar ratio of accumulated $CH_4$ and $CO_2$. (B) Molar ratio of accumulated $CH_4$ and total organic matter. (C) Molar ratio of accumulated $CO_2$ and total organic matter.

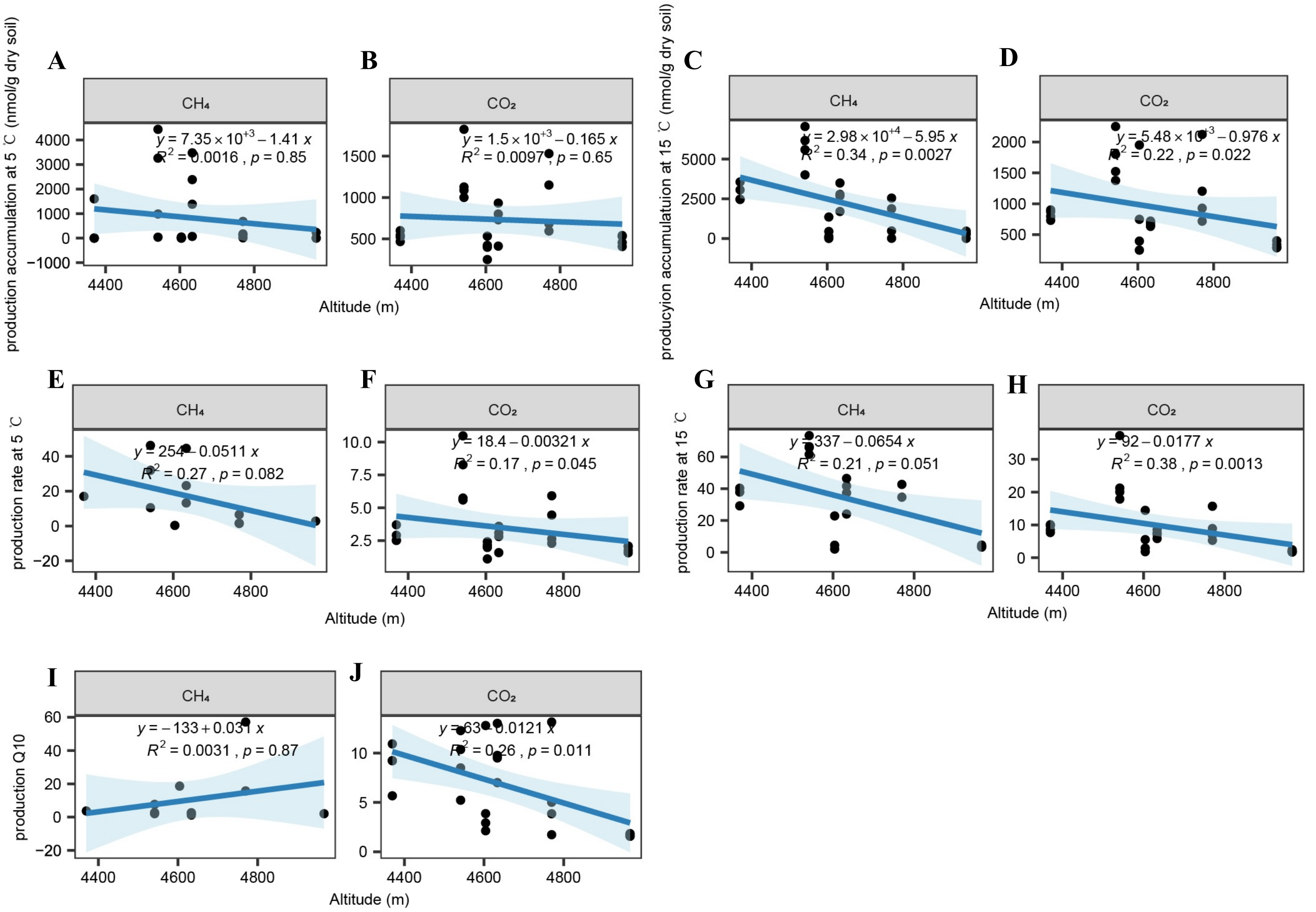


**Fig. S5. Correlation of the gas production rate and Q10 with altitude.** (A) and (B) Correlation of accumulated $CH_4$ and $CO_2$ with altitude for samples incubated at 5°C, respectively. (C) and (D) Correlation of accumulated $CH_4$ and $CO_2$ with altitude for samples incubated at 15°C, respectively. (E) and (F) Correlation of the $CH_4$ and $CO_2$ production rate with altitude for samples incubated at 5°C, respectively. (G) and (H) Correlation of the $CH_4$ and $CO_2$ production rate with altitude for samples incubated at 15°C, respectively. (I) and (J) Correlation of Q10 of the $CH_4$ and $CO_2$ with altitude, respectively.

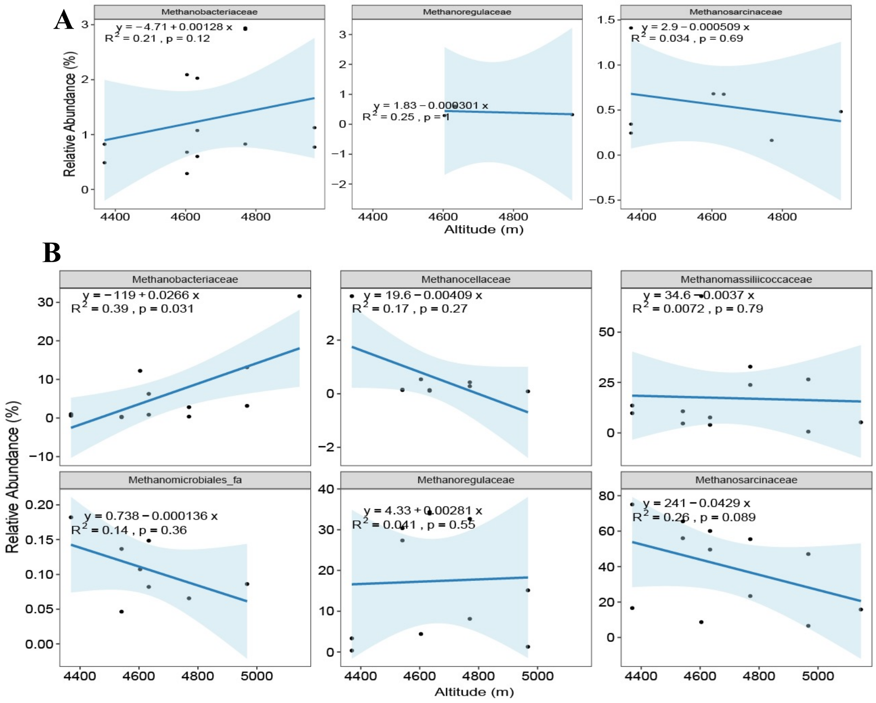


**Fig. S6. Relative abundance of major archaeal families and their correlations with altitude. (a)** correlation of methanogens family in original samples with altitude (b) correlation of methanogens family in samples after incubation at 5°C with altitude.

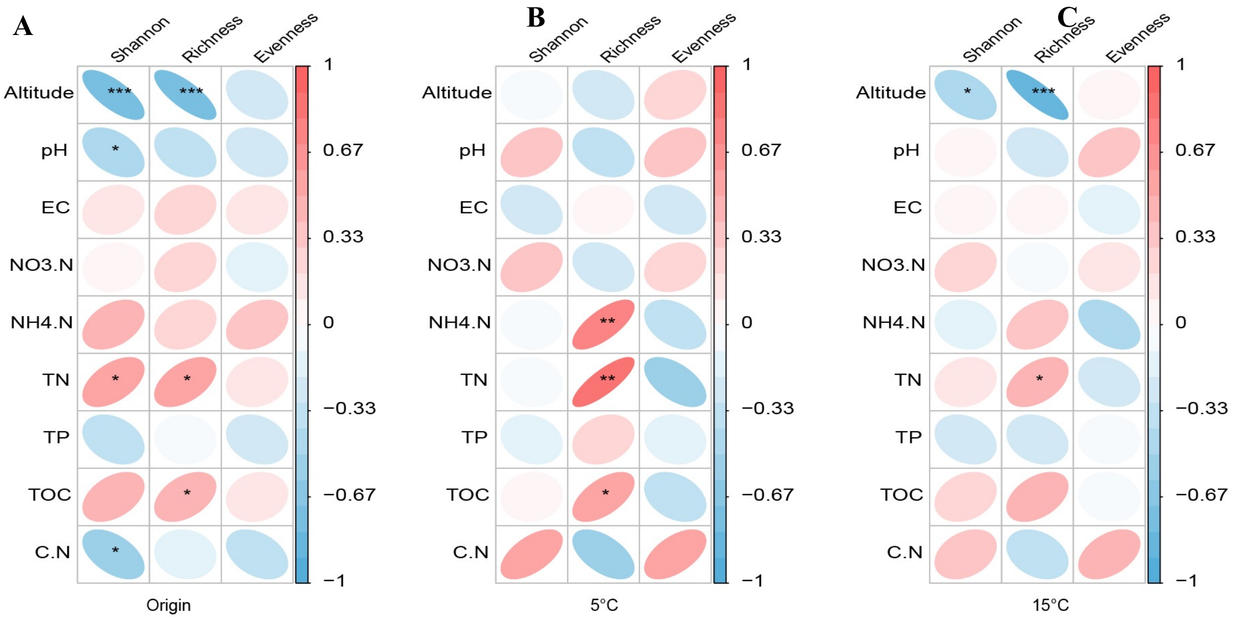


**Fig. S7. Factors influence alpha diversity of archaeal communities in the original (A), post-incubation samples at 5°C (B) and 15°C (C), respectively.**

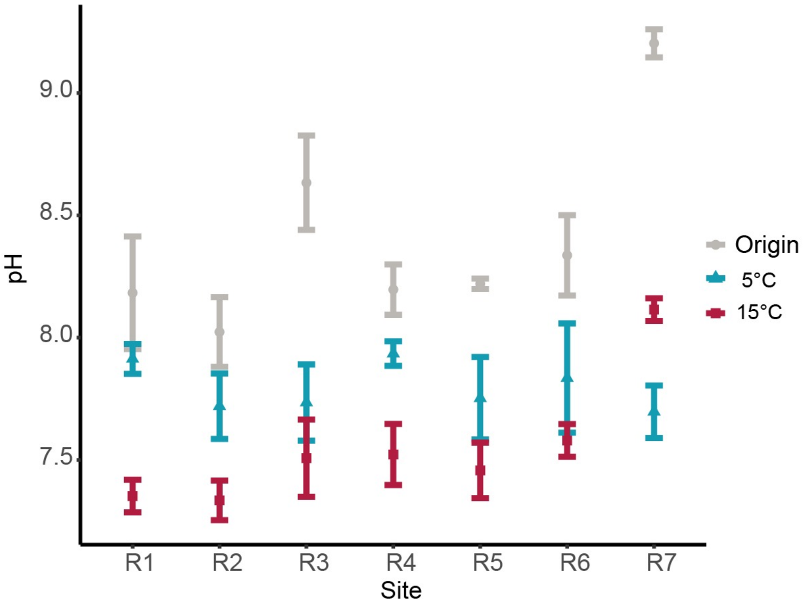


**Fig. S8. pH before and after incubation.**

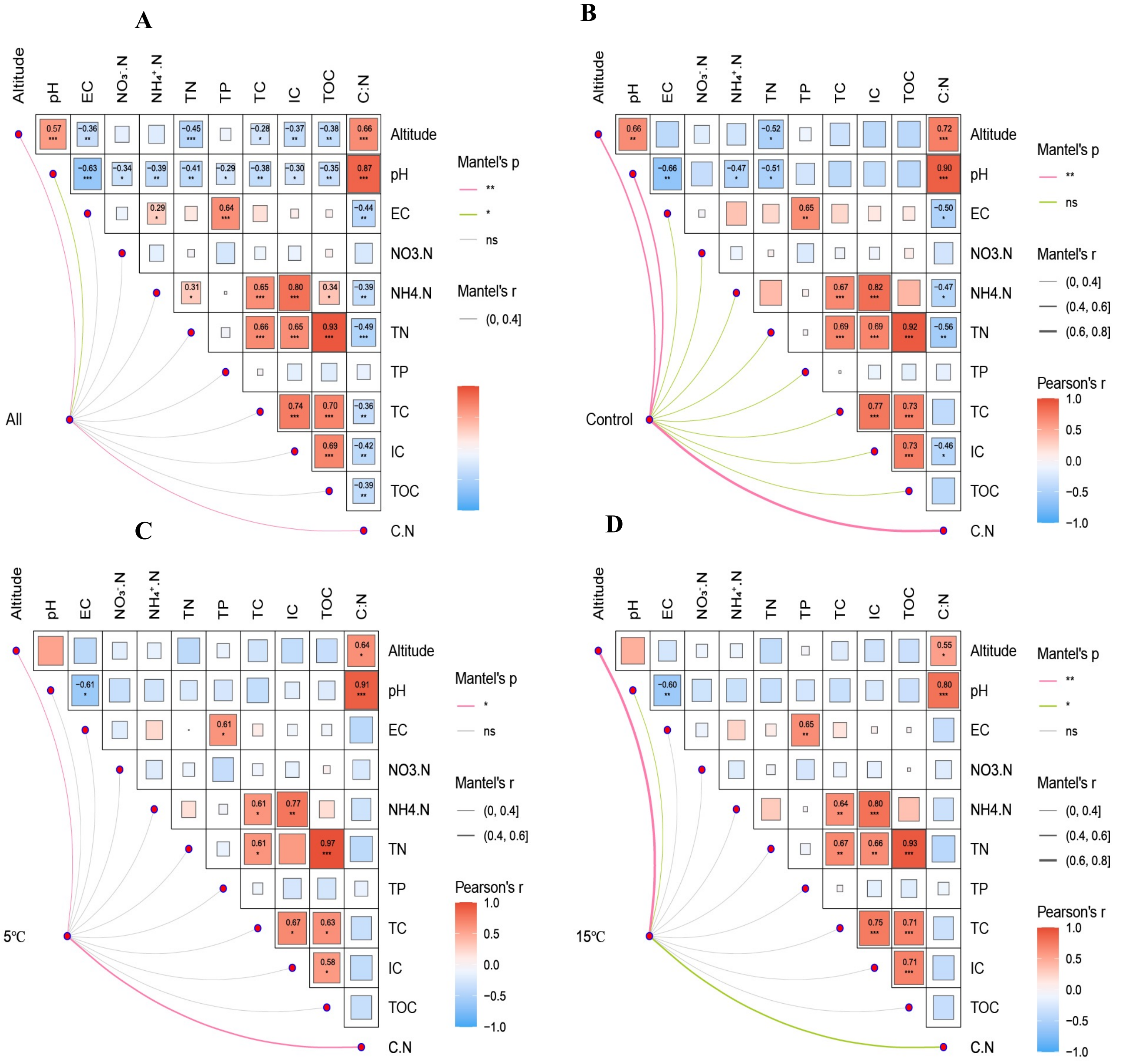


**Fig. S9. Factors that influence archaea communities.** Mantel test of factors with archaeal community of (A) all , (B) original samples, (C) archaeal communities at 5°C and (D) at 15°C.

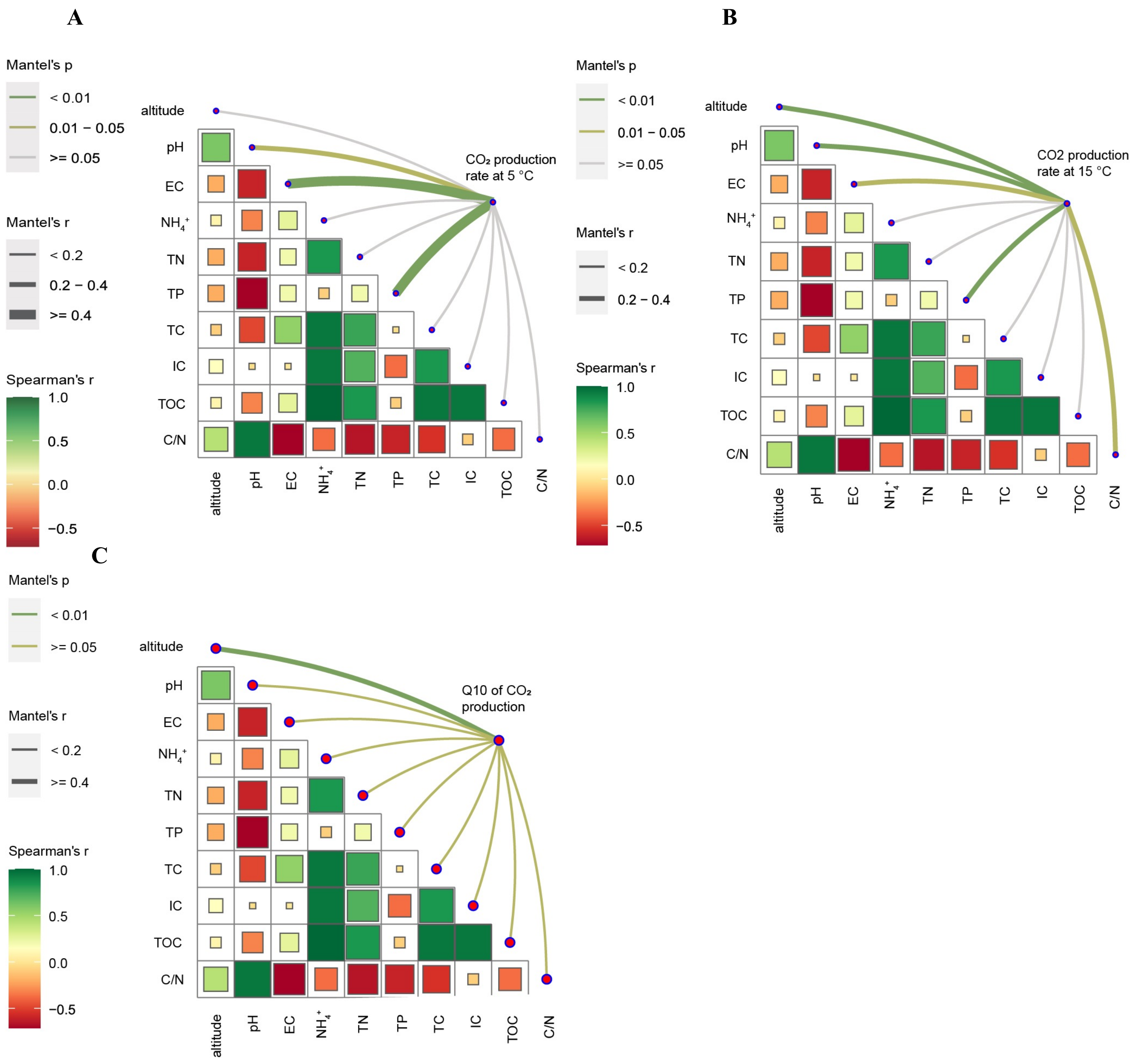


**Fig. S10. Factors influence $CO_2$ production rate, and Q10.** (A) Mantel test showing the relationship between sediment properties and $CO_2$ production rate at 5°C. (B) Mantel test showing the relationship between sediment properties and total $CO_2$ accumulation at 15°C. (C) Mantel test showing the relationship between sediment properties and Q10 of $CO_2$ production.

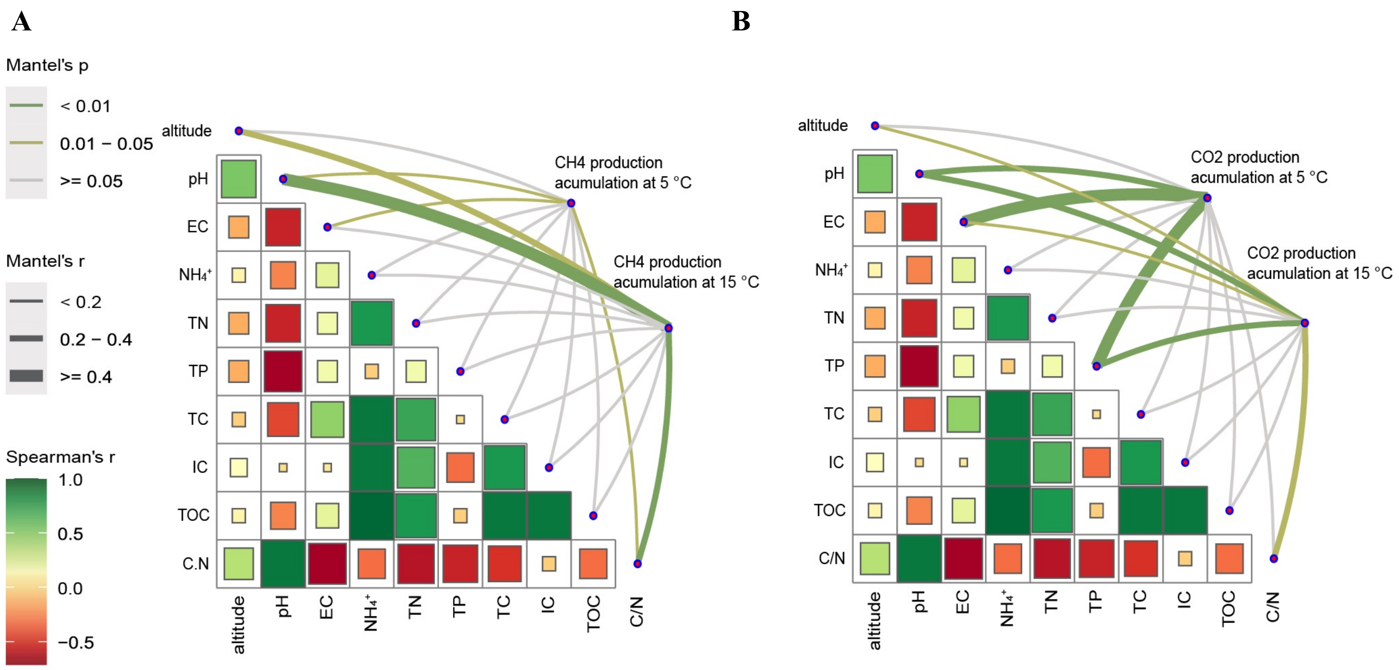


**Fig. S11. Factors influence $CH_4$ and $CO_2$ accumulation.** (A) Mantel test showing the relationship between sediment properties and accumulated $CH_4$ at 5°C and 15°C. (B) Mantel test showing the relationship between sediment properties and total $CO_2$ accumulation at 5°C and 15°C.
